# Generalized Hypergeometric Distributions Generated by Birth-Death Process in Bioinformatics

**DOI:** 10.1101/2022.02.02.478865

**Authors:** Vladimir A. Kuznetsov, Andre Grageda, Davood Farbod

## Abstract

Modern high-throughput biological systems detection methods generate empirical frequency distributions (EFD) which exhibit complex forms and have long right-side tails. Such EFD are often observed in normal and pathological processes, of which the probabilistic properties are essential, but the underlying probability mechanisms are poorly understood. To better understand the probability mechanisms driving biological complexity and the pathological role of extreme values, we propose that the observed skewed discrete distributions are generated by non-linear transition rates of birth and death processes (BDPs). We introduce a (3d+1)-parameter Generalized Gaussian Hypergeometric Probability ((3d+1)-GHP) model with the probabilities defined by a stationary solution of generalized BDP (g-BDP) and represented by generalized hypergeometric series with regularly varying function properties. We study the Regularly Varying 3d-Parameter Generalized Gaussian Hypergeometric Probability (3d-RGHP) function’s regular variation properties, asymptotically constant slow varying component, unimodality and upward/ downward convexity which allows us to specify a family of 3d-RGHP models and study their analytical and numerical characteristics. The frequency distribution of unique mutations occurring in the human genome of patients with melanoma have been analyzed as an example application of our theory in bioinformatics. The results show that the parameterized model not only fits the ‘heavy tail’ well, but also the entire EFD taken on the complete experimental outcome space. Our model provides a rigorous and flexible mathematical framework for analysis and application of skewed distributions generated by BDPs which often occur in bioinformatics and big data science.

## 1 Introduction

Today, high-throughput biological systems detection methods and bioinformatics generate enormous amounts of information (’big data’) about characteristics of large-scale molecular and cellular random processes which drive the evolution, interaction, complexity and integrity of such systems. Such complex data sets are subject to statistical bioinformatics and mathematical modeling that analyze and extract data to generate knowledge. These information science disciplines change biology, other fields, and introduce new technological challenges. Development of big data in bioinformatics assist in solving fundamental challenges in biomedical disciplines and presents unimaginable prospects in data technology. Accordingly, novel complex and practically useful mathematical models need to be developed and applied.

The theories of probability and stochastic processes play important roles in the modeling and analysis of big data in bioinformatics. It has been observed that at the system level, molecular and cellular characteristics often range from zero to 10^4^ – 10^7^ and their empirical frequency distribution (EFD) often exhibits skewed long-right side tail [24, 25, 26, 29] which is also over-dispersed [25, 26, 29]. The diverse EFD forms are observed in the normal and pathological processes, that probabilistic properties are essential, but underlie probability mechanisms are poorly understood.

The Birth-Death Process (BDP) with different functional forms of birth and death rates as the continuous-time and discrete states random process is one of the biologically reasonable models frequently used in bioinformatics and large-scale biomolecular data analysis. Importantly, the stationary distributions that are defined by the BDP could exhibit a skewed form having a power law-like right-side tail [25, 26, 27]. We have used linear BDP to model the gene expression level statistics that identified the skewed stationary probability distribution called Kolmogorov-Waring Distribution (KWD) and applied this model to data analysis of the Cancer Genome Anatomy Project (CGAP, NIH) which generated hundreds of sequencing and microarray data sets. The KWD model has been successfully quantified a nd a pplied to the analysis of Serial Analysis of Gene Expression (SAGE) and microarray data sets collected by CGAP and led us to the discovery of the power law-like frequency distribution at the gene expression level in the transcriptome of different bulk tissue samples (including cancer and normal tissues), stem cells, cell lines of many eukaryotes. The KWD not only proper fits well to such skewed EFDs with power law-like fat tail, but also has explained observed systematic changes of the frequency distribution form as a function of sample size and biomedical conditions.

Currently, there are many successful applications of BDP generated skewed frequency distribution models in diverse fields of bioinformatics and systems biology. For example, in an analysis of protein domain occurrences in proteins of a proteome of organisms [25, 26, 28, 34], the number of expressed genes in the transcribed genome of cells [25, 26, 28], the DNA-protein binding avidity of the transcription factors (TFs) - proteins bound the specific DNA sites (BS) and regulate gene transcription [29, 32, 40], the number of R-loops formation sequences per gene region in the genome of an organism [41], the number of mutations in a gene of cancer cells [35].

Using skewed distributions according to the variety and diversity of bioinformatics data, two- and more parameter distribution models with linear birth and death rates as a function of the state of the process have been developed and used in big data analysis. Two-parameter Waring distribution [25, 26], three-parameter KWD [28, 30, 31], regularly varying hypergeometric frequency distributions with two, three and more parameters [2, 3, 4, 5, 6, 11, 13, 15] have been developed and applied to the analysis of the EFD generated by bioinformatics data sets.

In bioinformatics data analysis the multi-parameter probability models based on the BDP have been also developed [3, 6, 12, 21, 26, 28, 29, 30, 31, 34]. Kuznetsov has developed the generalized KWD (g-KWD) [26, 28, 29, 30, 31], applications of BDP with several parameters to protein domains evolution in proteins could be found in [21, 34], a spatial subclass of “multi-parameter regularly varying generalized hypergeometric distribution of the second type” has been introduced by Danielian et al.[12] whose considered the skewed distributions generated by BDP via analysis of bioinformatics data.

We specifically refer to the pioneer studies by Kemp [22] and Maki [33] who have used stationary Probability Distribution Function (s-PDF) generated by BDP to develop a basic theory of the multi-parameter distributions. Both scientists have considered the rational functions to model transition rates of integer argument *x* ϵ {0, 1, 2, …} [22, 33]; see [19, 29] for more references.

Referring to the previous studies [3, 6, 26, 28, 30, 31] Danielian et al.[12] have applied a regular variation theory to the skewed distributions generated by the BDP occurring via analysis of bioinformatics data, introduced multi-parameter regularly varying generalized hypergeometric distribution of the second type. This theory specified some mathematical properties of the EFD, called *statistical (empirical facts*, which have been observed in bioinformatics data sets and are systematically reproducible in biomolecular systems. The facts follow skewness to the right, regularly varying at infinity, unimodality, stability by estimated parameter values, and convexity. These properties are also commonly observed in many other evolving complex systems. It was proposed that if a PDF (or probability law) satisfies these essential properties, then the PDF could properly be a mathematical framework for bioinformatics applications and statistical data analysis [3, 6]. However, more systematic, rigorous theory and data-driven applications need to be developed.

To better understand probabilistic mechanisms driving complexity and pathological role of extreme values in biosystems, we hypothesized that the observed skewed distribution of random event(s) varying at a large scale in these systems is a result of non-linear and mutually independent transition rate functions defined b y several probabilistic mechanisms. We propose that the asymptotic theory of the generalized skewed distributions with extreme value tail generated by BDP provide a rigorous mathematical framework in a search for plausible probabilistic mechanisms essential for the biosystem’s evolution and pathological regime. Our motivation is to select and develop mathematically adequate family of non-linear probability distributions by verifying the *statistical facts*, offer a family of the models in particular the stationary probability distributions with more than two parameters and non-linear transition rate functions of state of the BDP. In this study, we propose a new *(3d+1)-Parameter Generalized Gaussian Hypergeometric Probability ((3d+1)-GHP)* distribution based on the hypergeometric series function and then provide some biological applications for it.

### Note 1.

Danielian et al.[12] have studied a practically meaningful subfamily of Generalized Hypergeometric Probability Distribution [19, 22] called by the authors the Generalized Hypergeometric Distribution of the Second Type. In this paper we consider a subfamily of (3d+1)-GHP as *Regularly Varying 3d-Parameter Generalized Gaussian Hypergeometric Probability Distribution (3d-RGHP)*.

The paper is formed as follows. In section 2 introduces the (3d+1)-GHP model as a stationary distribution of generalized BDP (g-BDP). In section 3 we prove the *statistical facts* for the subfamily 3d-RGHP as a model proper for bioinformatics data sets and biomedical systems. Section 4 presents numerical examples of PDF of the 3d-RGHP, and an analysis of the PDF in the log-log plot. Section 5 provides results of the (3d+1)-GHP and 3d-RGHP fitting to the EFD of unique mutations that occurred in melanoma genes genome-wide. Section 6 provides a discussion and conclusion.

## 2 Generalized BDP with (3d+1-GHP

Suppose that a random process is a family **X** = {*X*(*t*), *t* ϵ *T*} of random variable values *X* = *i*, *i* = {0, 1, 2, ..} such that the process is a Markov process with continuous time *t* ϵ *T* ≥ 0 and countable number of states *X* = *i* where *i* ϵ *S* = {0, 1, 2, …}.

Continuous time BDP is a practically applicable subset of the Markov chains models [16, 23]. The BDP {*X*(*t*), *t* ≥ 0} is described as a random time movement of the object between neighbor states with non-negative and finite transition rates. BDP model allows change state only for one step for short time interval (*t*, *t* + *δt*]: *E*_*i*_ → *E*_*i*+1_ as well as *E*_*i*_ → *E*_*i*–1_ if *i* ≥ 1. If *i* = 0, the theory allows *E*_0_ → *E*_1_. Assume that if the process at time *t* is in *E*_*i*_, then during (*t*, *t* + *δt*] the transitions *E*_*i*_ → *E*_*i*+1_ have probability *λ*_*i*_(*t*)*δt* + *o*(*δt*), *E*_*i*_ → *E*_*i*–1_ have probability *μ*_*i*_(*t*) + *o*(*δt*), probability no change *E*_*i*_ in state *i* be 1 – (*λ*_*i*_(*t*) + *μ*_*i*_(*t*))*δt* + *o*(*δt*) and a probability of more than one state change occurs equals *o*(*δt*). *o*(*h*) means that 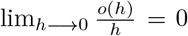, called negligible with respect to *h*.

Let {*p*_*i*_(*t*) = *P* (*X*(*t*) = *i*), where *i* ϵ *S* = {*i* = 0, 1, 2…}, denote the probability mass function (PMF) at time *t* ϵ [0, ∞) generated by the stochastic process {*X*(*t*), *t* ≥ 0}. According to the theory, at each infinitesimal time interval (*t, t* + *δt*] where *δt* → 0 the transition functions are differentiable functions and we can calculate birth and death transition rates

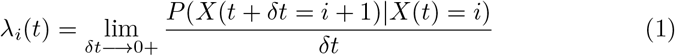

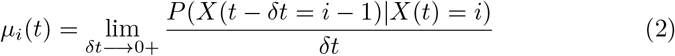

where ∞ > λ_*i*_(*t*) > 0 for *i* ≥ 0, ∞ > *μ*_*i*_(*t*) > 0, *i* ≥ 1, ∞ > *μ*_0_(*t*) ≥ 0.

In this section, we generalize the BDP model, considering the transition rates as non-linear multi-parameter functions of states. According to basic assumptions of the BDP model, when at time *t* a state of a population is *i*, the probability of a single birth occurring during the short time interval (*t, δt*) is defined by the conditional probabilities (1-2) which does not depend on time interval *δt* and for also at *δt* → 0

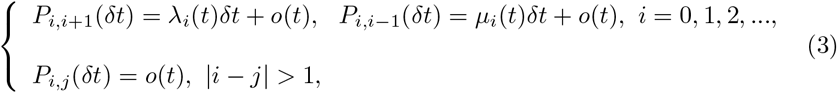

and we conclude that

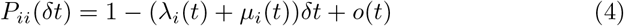

as *δt* → 0 and

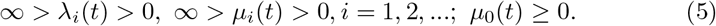

Using (1-2) and (3-4) and assuming a differentiation property of the probability functions *p*_*i*_(*t*), *i* = 0, 1, 2, …, the total changes of each probability in the BDPs, we have Kolmogorov’s BDP differential difference equations at *δt* → 0 [16], [23]:

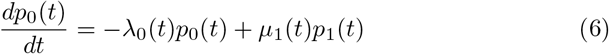

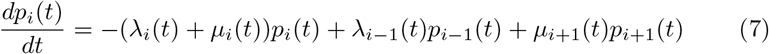

where *i* = 1, 2, … We also assume that

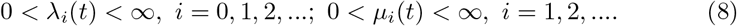

The BDP defined by (6-7) at any time is the following conditions

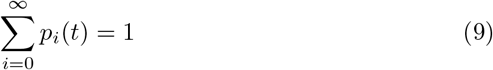

and other two Kolmogorov’s axioms [23, 37]. Also, according to Markov chain property the probability measure of the BDP is a right-continuous step-function, suggesting its ‘regularity’ property.

Additional specification of the properties of the BDP allows us to define the distribution laws with potential applications.

Let us consider long-time regimes of the BDP related to stationary and limiting characteristics and the corresponding probability functions [9, 16].

Let *n*(*t*_1_, *t*_2_) be a random number of events of the process within time interval [*t*_1_, *t*_2_] at time *s* ϵ [0, *t*]. The BDP is called stationary (or time-homogeneous) if the probability distributions of r.v.’s *n*(*t*_1_, *t*_2_) and *n*(*t*_1_ + *s*, *t*_2_ + *s*) are coincident *P* (*n*(*s, s* + *t*) = *i*) = *P* (*n*(0, *t*) = *i*) where *i* = 1, 2, … for *s* ϵ [0, *t*]. We will consider the stationary process BDP with non-zero limiting distribution. This proposes that all transition rate functions of the process are positive and there are no adsorption states, in particular we put *μ*_0_(*t*) = 0.

Let 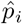, (*i* = 0,1,2, …) be a probability of stationary process in state *i* and 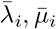 be transition rates of the process in this state.

#### Definition 1.

A PDF of random variable *X* = *i, i* ϵ *S* = {0, 1, 2, …} is a s-PDF of the process {*X*(*t*), *t* ϵ [0, ∞) given by (6-7) if the probabilities 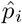, (*i* = 0, 1, 2, …) are defined by the non-trivial solution the system of algebraic equations

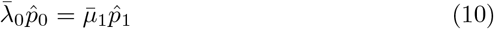

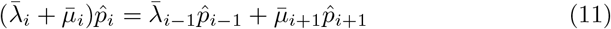

where *i* = 1, 2, …. These equations provide a singular point solution of the system of differential-deference BDP equations at

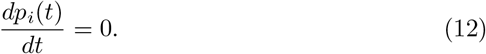

Note that probabilities of *X* of PDF is a discrete distribution function which also called a PMF.

Thus, we have

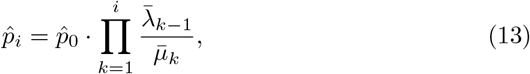

and

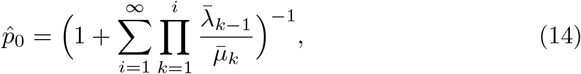

where 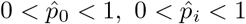, *i* = 1, 2, …

Note that

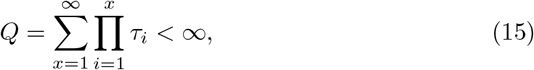

where *x* = 1, 2, …, 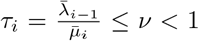 provides well-known necessary and sufficient conditions for the existence of the non-trivial stationary solution Eqs.(13-14).

To characterize asymptotic properties of the probability distribution in time, the time of the random process after its initiation can be discriminated on two intervals: [0, *t*_*n*_) called time of the transition phase (non-stationary process phase) and next time interval [*t*_*n*_, ∞), which refers to the long-time behavior, is called limiting or steady state phase.

We consider long-time asymptotic solution of BDP (6-7) allows determining this solution using

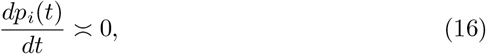

at *t* → ∞ in states *i* = 0, 1, 2, … The solution is the PMF that is also named steady-state.

The probabilities of PMF in the steady-states we denote by 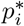, *i* = 0, 1, 2, , …..

In this study we assume that in the lim_*t*→∞_ *p*_0_(*t*) > 0.

Let us define the non-zero limiting probability function of (6-7) which takes r.v. *X*(*t*) at *t* → ∞:

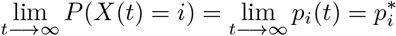

Let *i* be an initial state and *j* be a limiting in time state of a trajectory given by BDP (6-7) where {*i, j*} ∈ *S* = {0, 1, 2, …,}, i ≠ *j*, stationary PMF exists and is defined by the probabilities 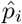, *i* ∈ *S* = {0, 1, 2, …,}.

### Definition 2.

The PMF is the limiting PMF (defined by probabilities 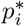, *i* = 0, 1, 2, …,) of the BDP if, for any transition from initial state *i* to a state *j*, the transition rates are defined by transition rates of the stationary distribution.

Based on the axioms of a probability and a definition of the Markov chain, we assume that in the 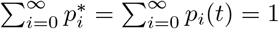, *t* ϵ *T* = [0, ∞), *i* ϵ *S* = {0, 1, 2…,}.

Note that the expression (15) is applicable for calculation of the limiting distribution after certain period of ‘transition phase’ of the process when starting from some *i* = *i*_*j*_ at time *t* > *t*_*c*_ > 0 the condition *τ* ≤ *ν*_*i*_ < 1 happen for all *i* > *i*_*j*_ (namely, in the right tail of the probability function).

The BDP {*X*(*t*), *t* ≥ 0} is an ergodic Markov chain if it is possible to go from every state to every state. This proposes that for any initial distribution of state probabilities *p*_*i*_(*t* = 0), *i* ϵ *S* = {0, 1, 2, …,} the transition rates are positive and the limiting probability distribution representing by 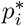 of random variable *i* exists due to stationary distribution exists. The conditions 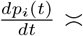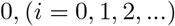 assume the asymptotic converge of *p*_*i*_(*t*) to 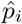 when *t* → ∞.

Thus, we have

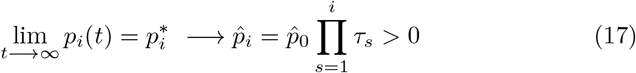

This suggests a long-time asymptotic converge of the BDP equations solutions to an isolated (stationary) solution.

Thus, the long-time limiting distribution is independent of the initial state while the stationary distribution is determined by the unique initial state distribution (it is a non-trivial singular solution of the BDP equations). The long-time limiting distribution is an asymptotic distribution, while the stationary distribution a special initial state distribution.

Now, having the above definitions and assumptions, we consider that the birth and death intensity rates between states may be approximated by polynomial or rational functions with integer argument *x* ϵ {0, 1, 2, …} [20, 22, 26, 28].

Let as consider a stationary BDP which birth and death rates given by quadratic functions of state *x* = {0, 1, 2, …} given with three positive parameters sets {*a, b, θ*_*λ*_} for birth rate and two parameters for death rate functions {*c*, *θ*_*μ*_} respectively. We consider these functions as the following

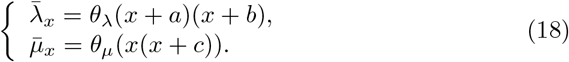

Below we will show that (18) allow to define the solution of stationary BDP in Gaussian hypergeometric series (GHS) terms [19, 17].

The GHS is a series 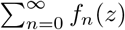 such that 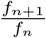 is a rational function of *n* [19, 17]. On factorization of the two polynomials in degree *n*, we obtain

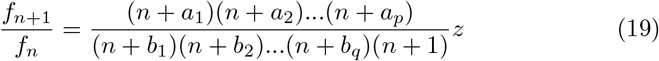

where *z* is the argument of the function, which may be equal to one (monic), or may be not. The factor (*n* + 1) may be result in factorization or may be not. Decomposition of (19) into rational form is depending on *p* + *q* constants and argument *z*.

For *p* = 2 and *q* = 1, *z* = *θ* where *θ* is a positive real value, we have the GHS

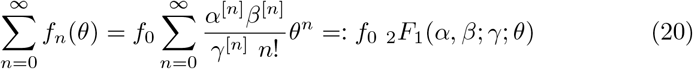

where *n* ϵ {0, 1, 2, …,}. The terms of series are characterized by parameters *α, β, γ* of Pochhammer’s symbols and series argument *θ*. Pochhammer’s symbol is standard ascending factorials notation *a*^[*n*]^ that denotes 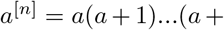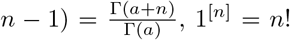. In the literature, the (20) are associated with the families of Generalized Hypergeometric Probability (GHP) [19, 22, 36].

It is straightforward to show that using the expression (13-14) and the birth and death rates given by (18), the probabilities 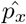 are converted to the terms power series function of an argument *θ* with exponent parameter *x* ϵ {0, 1, 2, …} and the coefficients defined as the rational functions given by *a*^[*x*]^, *b*^[*x*]^ and (*c* + 1)^[*x*]^

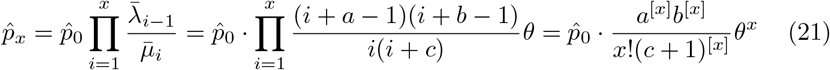

where 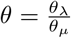 and 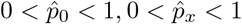, *x* = 1, 2, ….

Finally, putting *a* = *α, b* = *β*, (*c* + 1) = *γ* in the (21), we obtain the probabilities *p*_*i*_, *i* ϵ *S* = {0, 1, 2, …} in the forms of GHS terms and 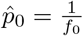

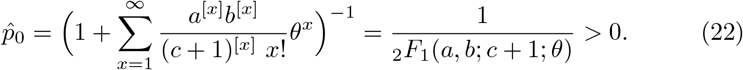

Note, at *b* = 1 or *a* = 1 the probabilities (21-22) are the probabilities of KWD with linear birth and death rate functions of *X* = *x* [25, 26], and if *b* ≠ 1 or *a* ≠ 1 the probabilities (21-22) could be considered as a specific case of g-KWD where the birth and death rate functions are polynomials of *x* with degree 2.

#### Corollary 1.

The probabilities 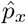, *x* = 0, 1, 2, … of s-BDP with transition rates (18) are defined by GHS terms and calculated by the following

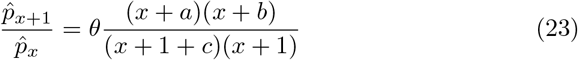

at 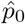 defined by (22).

At 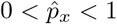, *x* = 1, 2, …, where the parameters (*a, b, c*) are defined by the properties of the birth and death rate functions (18), (0 < *a* < ∞, 0 < *b* < ∞, 0 < *c* < ∞). Referring to (23) and due to 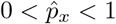, *x* = 0, 1, 2, …,

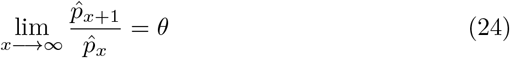

where the values of the parameter *θ* should be positive and limited by interval (0, 1].

In this study we consider a generalization of BDP with birth and death rate functions defined as a finite product of the functions given by quadratic birth and death rate functions of *X* = *x* in form (18) which parameters may have different non-negative values. We introduce the generalized form of the transition rate functions as follows

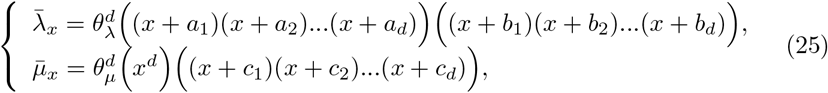

where 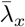 and 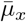 are the birth and death functions of random variable *X* = *x* respectively, where *x* ϵ *S* = {1, 2, …, *x* …} and *X* the is a countable random variable. *a*_*j*_, *b*_*j*_, *c*_*j*_ are the parameters 0 < *a*_*j*_ < ∞, 0 < *b*_*j*_ < ∞, 0 < *c*_*j*_ < ∞ where *j* = 1, 2, …, *d* are the elements of the 3d-parameter set *V* = {*a*_*j*_, *b*_*j*_, *c*_*j*_}. *θ*_*λ*_ = *const* > 0, *θ*_*μ*_ = *const* > 0; 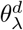 and 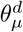 are the highest powers of *x* in the polynomials with degree *d*. Let denote 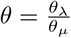.

Using (13-14) and (25) we can present the probabilities 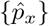 with (3d + 1) unknown parameters of the birth and death rate functions as follows

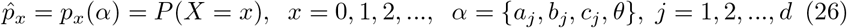

takes the following form

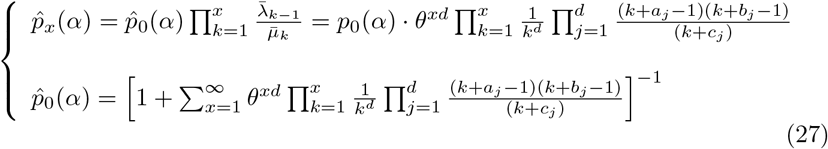

where *x* ϵ *S* = {1, 2, …, *x*, …}. *α* = {*a*_*j*_, *b*_*j*_, *c*_*j*_, *θ*} is a set the unknown parameters such that 0 < *a*_*j*_ < ∞, 0 < *b*_*j*_ < ∞, 0 < *c*_*j*_ < ∞, where *j* = 1, 2, …, *d*, 0 < *θ* ≤ 1. In general case, *d* could be an unknown parameter of the model.

Note, 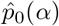 is the probability normalization factor providing 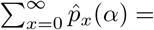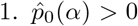 provides a condition of an existence of the s-PDF. At 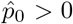 the stationary distribution existence is well-defined by convergence of Gaussian hy-pergeometric series. 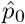 is a function of all parameters and could be interpreted as a probability of non-observed events in experiment outcome space, that is defined by parameters of the distribution.

Having (27), we can obtain the stationary probability measure *F*_*x*_ (*α*) = *P* (*X* ≤ *x, α*) (also called as the distribution of random variable *X*)

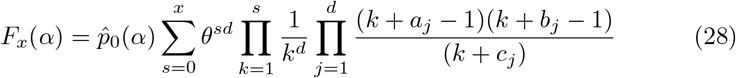

Using Pochhammer’s symbol we obtain (27) in the form

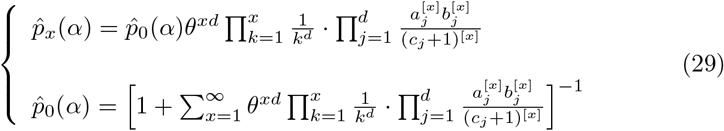

Note, 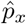, *x* = 0, 1, 2, … in (27) (as well as (29)) could be presented by generalized hypergeometric series [17, 19].

This follows from recursive form of state probabilities (27) as

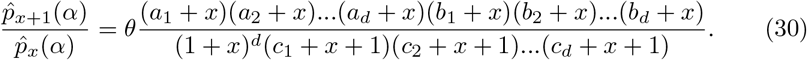

### Corollary 2.

The g-BDP defined by birth and death rate functions given by (25) generates the stationary probabilities (27) and a probability measure (28) called here the (3d+1)-GHP.

The 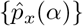 at 0, *d* = 1, 2, …, provides a broad family of s-PDF (27) with the functionally diverse properties of the birth and death transition rate functions (25).

Note that (30) could be used to study asymptotic properties of the (3d+1)-GHP, specifically in the case when both birth and death rate functions are growing when an argument increases *x* → ∞.

In this case

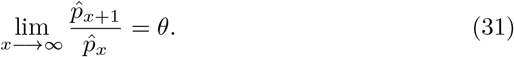

### Corollary 3.

Let us consider (30). If *θ* > 1 then a stationary BDP process not exists; if 0 < *θ* < 1 then the PMF tail is unimodal, strictly decreases and the tail becomes longer if *θ* → 1–; if *θ* = 1 the existence and asymptotic properties of the stationary distribution is defined by the relationships of the parameters {*a*_*j*_, *b*_*j*_, *c*_*j*_}, *j* = 1, 2, …, *d* (see the next section).

Let *x* ∈ *S* = {1, 2, …} and consider an asymptotic property of birth and death rate functions defined by (25) at *x* → ∞. We have

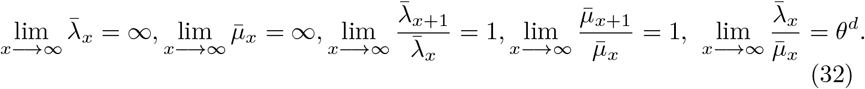

The last expression is most interesting because it shows that when the rates of the both BDPs increase at *x* → ∞ their ratio is positive constant and if the 0 < *θ* < 1 than the rate of death will be larger than the rate of birth. Thus, the stationary BDP exists; furthermore, the PMF tail becomes shorter if *d* be larger.

Let propose that PMF given by (27) exists and PMF unique at *θ* = 1. In this case 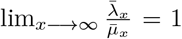 and at the fixed other parameters the PMF tail be the longest. Thus, our model gives us a characterization of ‘moderate growth’ for birth and death transition functions at asymptotic increasing the of *X* = *x*. Our previous studies have shown that a parameterization of the PFDs using skewed EFDs often provides *θ* = 1 – *ϵ*, 0 ≤ *ϵ* ≪ 1, referring to the case of most long tail of PDFs.

Now, we can consider the (27) to identify the relationships between parameters of the transition rate functions satisfying unimodality, convexity, regularity, skewness - the important mathematical characteristics commonly observed in large scale data sets of bioinformatics. We will to consider several basic asymptotic properties of our model in the case *θ* = 1.

## 3 Extracting a regularly varying distribution

Let us have the following definitions.

#### Definition 3

([12]). The distribution *p*_*x*_(*α*) varies regularly at infinity with exponent (−*ρ*) if it can be stated as

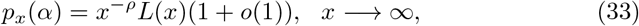

where *L*(*x*) > 0 for *x* = 1, 2, …, and for *s* = 2, 3, …, 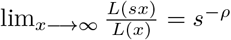.

#### Definition 4

([12]). Assume that, for *s* = 2, 3, …, the limit exists

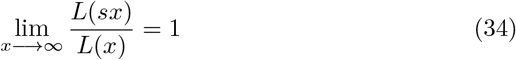

then we say *p*_*x*_(*α*) shows the asymptotically constant slowly varying component if in the formula (34)

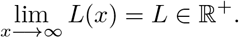

Now, we propose the following theorem.

#### Theorem 1.

*The model (3d+1)-GHP* (27) varies regularly at infinity if and only if *θ* = 1 and the exponent of the regular variation of 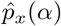 is equal to

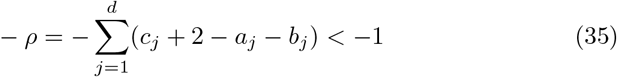

Meanwhile, 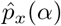 shows the asymptotically constant slowly varying component as

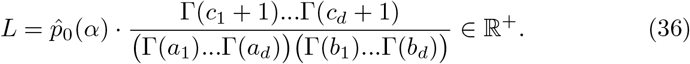

*Proof.* By using formula

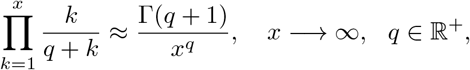

we obtain an approximation as follows (for *j* = 1, 2, …, *d*):

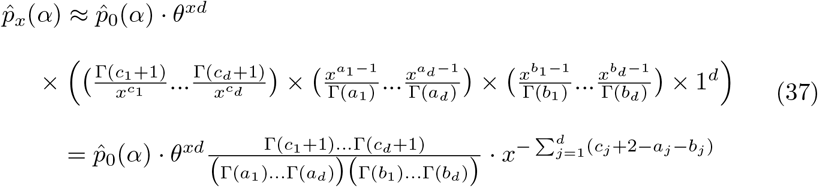

when *x* → ∞ and based on (33) we have 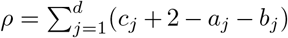 such that (−*ρ*) in (35) is the exponent of regular variation of 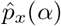. From (37) we turns out 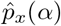 in (27) varies regularly at infinity if and only if *θ* = 1.

Additionally, by a limit *L* = lim_*x*→∞_*L*(*x*), we consider asymptotically constant slowly varying component (if exits). From (37) we see that *L* presented in (36) is an asymptotically constant slowly varying component for 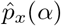. The Theorem 1 is proved.

### Remark 1.

Let, in the (3d+1)-GHP model (27), *θ* = 1. Then, the exponent 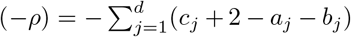 of regular variation holds the condition *ρ* > 1.

Hence, now based on the Theorem 1 and from (3d+1)-GHP distribution family (27) we can extract 3d-RGHP model with the following PMF:

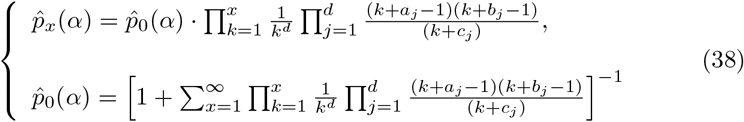

where 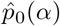 is the normalization factor, *x* = 0, 1, 2, …, *α* = (*a*_*j*_, *b*_*j*_, *c*_*j*_), *j* = 1, 2, …, *d*, is the unknown parameter such that 0 < *a*_*j*_ < ∞, 0 < *b*_*j*_ < ∞, 0 < *c*_*j*_ < ∞, *j* = 1, …, *d*. Also *a*_1_, …, *a*_*d*_ and *b*_1_, …, *b*_*d*_ are called numerator parameters and *c*_1_, …, *c*_*d*_ denominator parameters. We call the model (38) as 3d-RGHP.

Having (28), the stationary probability measure *F*_*x*_ (*α*) = *P* (*X* ≤ *x, α*) at *θ* = 1 is the following

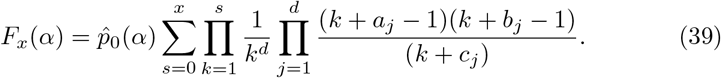

We call (39) as 3d-RGHP. Its PMF is written in the form (38)

### Remark 2.

Assume that, in the PMF (38) *d* = 1. Then it receives the three-parameter regularly varying generalized hypergeometric frequency distribution as a special case (see [11, 14]).

### 3.1 The log-log plot of 3d-RGHP

In this subsection versus 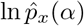 is considered. Let us establish the following theorem.

#### Theorem 2.

The 3d-RGHP model (38) is upward/downward convexity.

*Proof.* From 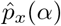 in (38) we have

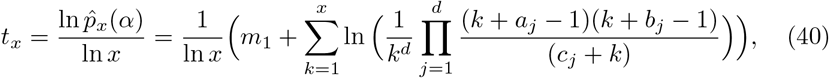

where *x* = 2, 3, …, and 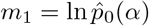.

Now, it can be written

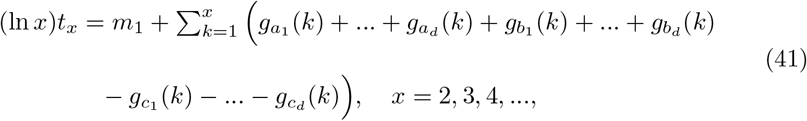

where for *ϵ* ϵ ℝ^+^ and *t* ϵ [1, ∞) and 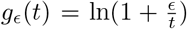. The function *g*_*ϵ*_ (*t*) is bounded and increasing. Compared to Danielian and Astola [11], it is readily seen that for *x* = 1, 2, …,

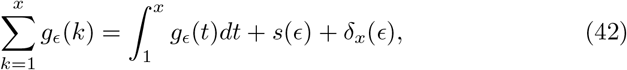

where *s*(*ϵ*) is constant and lim_*x*→∞_*δ*_*x*_(*ϵ*) = 0.

If we set

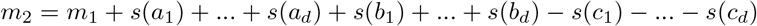

then by substituting (42) into (41) we get the following when *x* → ∞

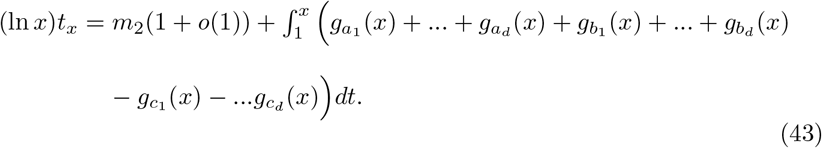

Moreover, compared to Danielian and Astola [11], for *ϵ* ∈ (0, ∞) and *x* = 1, 2, …,

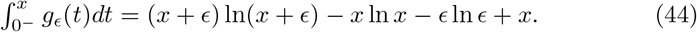

By substituting (44) into (43) it can be obtained when *x* → ∞

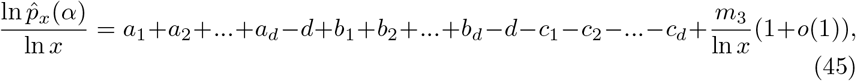

(*m*_3_ is some real constant). By (45) it follows that, at least for large values of *x*, the deviations of 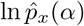 versus ln *x* from the straight line

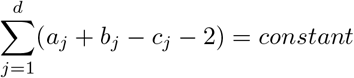

are small which shows upward/downward convexity. The proof of Theorem 2 is finished.

### 3.2 Convexity and unimodality

Unimodality of the 3d-RGHP and convexity on the right side PDF tail are essential features for frequency distributions in bioinformatics data sets. For details about this see [3, 6, 10, 21, 24, 25, 26, 28, 29, 30, 32, 34]. So, in this subsection, we shall attempt to prove such features for the 3d-RGHP model. We give a theorem as follows.

#### Theorem 3.

The 3d-RGHP model (38) is unimodal and asymptotically downward convex.

*Proof.* From 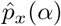 in (38) we obtain recursive formula for *x* = 0, 1, 2, …, as follows

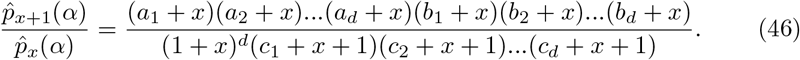

Let us rewrite the recursive (46) as

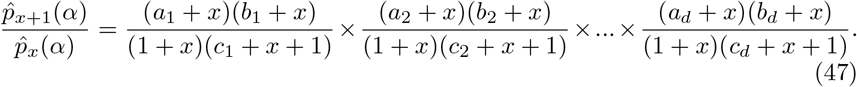

We investigate unimodality of 3d-RGHP 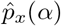 for some cases.

*Case 1.* Assume that, in (47) (in its first expression of right side), *min*(*a*_1_, *b*_1_) < 1. For example if *a*_1_ < 1 then 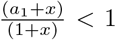 and increases when *x* increases. In view of *b*_1_ < *c*_1_ + 1, the expression 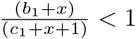 and increases when *x* increases. So, their product 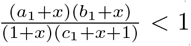 and increases when *x* increases. Similarly, this situation can be verified for 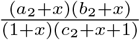 and 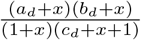. Therefore, based on (47)

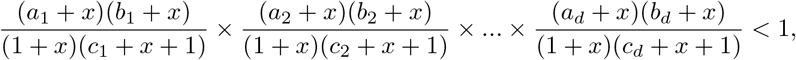

and the model 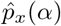 decreases and is downward convex. This concludes that 3d-RGHP model 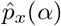 is unimodal.

*Case 2.* Suppose that *w*_*j*_ = (*c*_*j*_ + 2 – *a*_*j*_ – *b*_*j*_), *j* = 1, 2, …, *d*. From (47) let us write

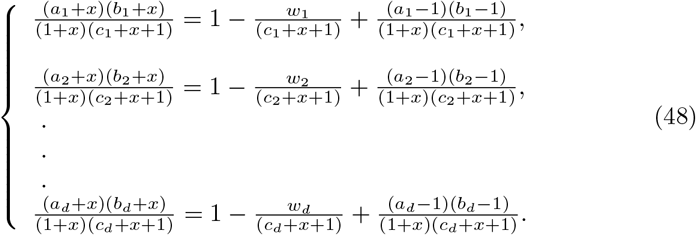

From (48), it can be concluded that if (*a*_*j*_ – 1)(*b*_*j*_ – 1) ≤ *w*_*j*_ then the Case 1 is received. Moreover, if (*a*_*j*_ – 1)(*b*_*j*_ – 1) > *w*_*j*_ then for enough large *x* the 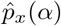 is again downward convex and in our cases is unimodal.

The proof of Theorem 3 is completed.

The following corollary is given.

#### Corollary 4.

Based on the Theorems 1, 2 and 3 we conclude that PMF and corresponding 3d-RGHP defined by probabilities (38) can be considered as a *new* non-linear and multi-parameter probabilistic model for the needs of data analysis and statistical inferences in bioinformatics and relevant fields.

## 4 Numerical analysis of the 3d-RGHP parameter: skewness variation and limitation of power law property

In Theorem 2, section 3, we proved that the deviations of 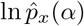 versus ln *x* from the straight line could be small, at least for some large values of *x*. However, the theory does not estimate how long the power law-like region on *x* should be, how soon it may occur, and how stable this extreme region may be when the parameter set is changed. To answer these questions, we fixed the value *d* = 2 and *θ* = 1 and use 4 distinct sets of parameters.

We considered 3d-RGHP in several special cases. We considered *θ* = 1 as the case corresponding to regularly varying function property. Furthermore, we set *d* = 2 and thus modeled the 6-RGHP for *α* = (*a*_1_, *a*_2_, *b*_1_, *b*_2_, *c*_1_, *c*_2_). We gradually increased all or some of the parameters. The values of *ρ* corresponding to the selected six-parameter sets were calculated. All values are given as follows

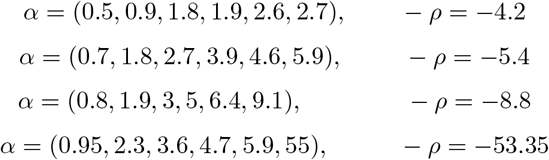

We plotted the PMF of 3d-RGHP (38) for the parameter sets at *d* = 2 (i.e. 6-RGHP). As proved in Theorem 3, Figures 1(A-D) show that for the different parameter sets the PMFs are unimodal (mode is observed at *x* = 1). However, if the gradual increasing all or some of the parameters, a curvature of the right-side tail of PMF was negatively corbeled fatness of the tail end. We also observed significant shift and variation of the power law-like right-side tail of the PMFs. Indeed, Figures 1(E-H) show the log-log plots of the double-truncated 3d-RGHP for the case 6-RGHP with different values of the parameters presented in Figures 1(A-D). We specified the cutoff values of random variable *x* interval that provide approximately similar slops of the function tails (asymptotic power law-like exponents). The left-side cutoffs a nd the right end of the intervals follows: In Figure 1-E, *x* ϵ [5, 15]; Figure 1-F, *x* ϵ [10, 20]; Figure 1-G, *x* ϵ [15, 25]; Figure 1-H, *x* ϵ [20, 30]. Figures 1(E-H) show that the power law tail interval is highly sensitive to parameter values, and a significant fraction of the probabilities within random variable intervals are significantly deviated from the straight line. Our numerical analysis observations are agreed with the variation of the value of regular variation *ρ*, at which is (negative) exponent of the PMF (at *θ* = 1).

**Figure 1:**
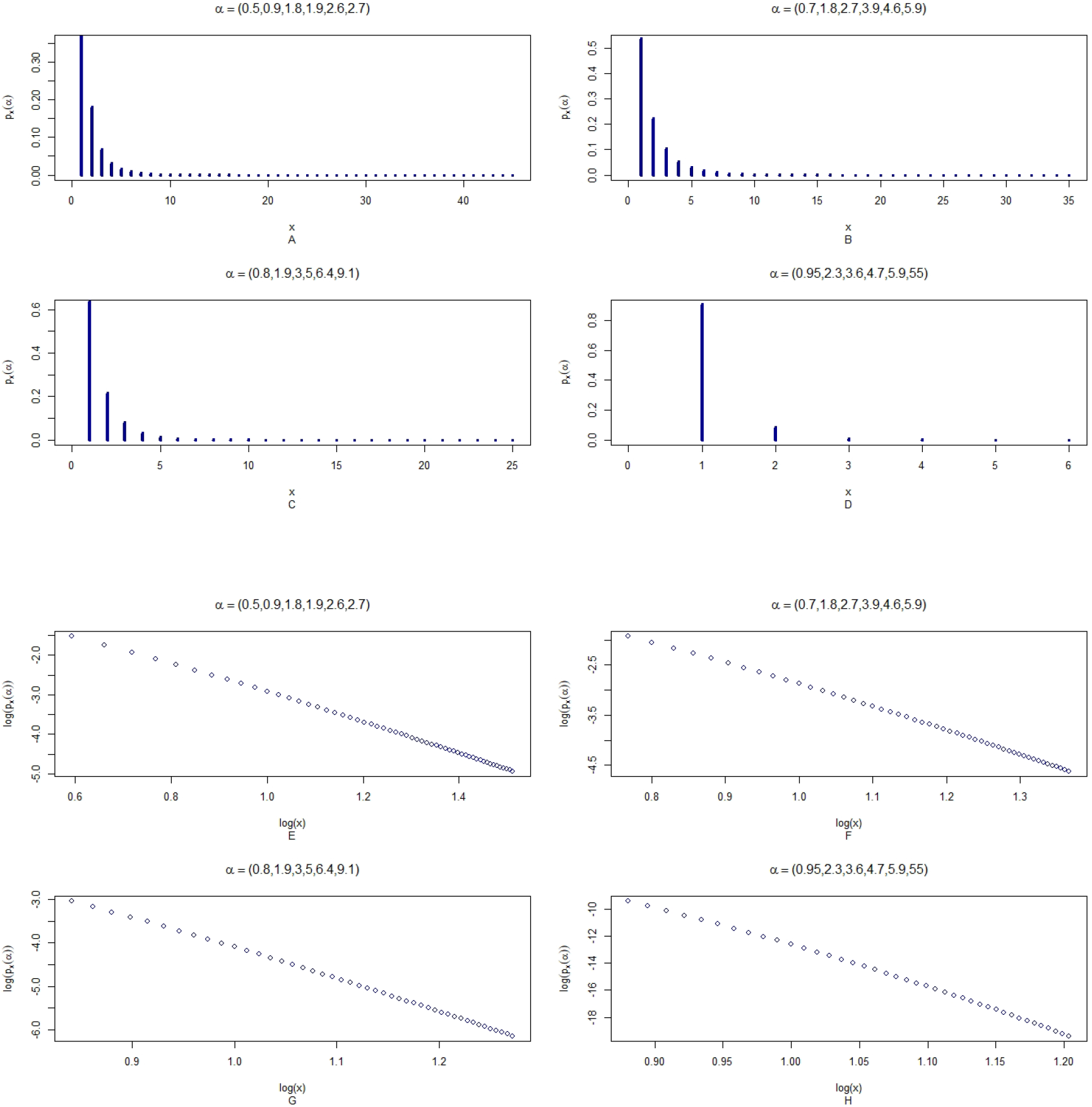
Illustrations of the PMF and log-log plots of 3d-RGHP model (38) when *d* = 2 (6-RGHP) for possible values of six parameters *α* = (*a*_1_, *a*_2_, *b*_1_, *b*_2_, *c*_1_, *c*_2_).

Our numerical analysis observations are agreed with the variation of the value of regular variation *ρ*, at which is (negative) exponent of the PMF (at *θ* = 1).

## 5 Application of the 3d-RGHP model. Distribution of the number of unique mutated samples in genes of melanoma patient’s genomes

To demonstrate the ability of our (3d+1)-GHP (27) and 3d-RGHP (38) models to analyze complex forms of the EFD with heavy tail on the right, we have downloaded somatic mutation data for the genes in several human cancers from the COSMIC database (https://cancer.sanger.ac.uk/cosmic). We studied the frequency distribution of the number of unique gene loci mutated samples detected in the large collection of human melanomas. Our data set used in this study, includes 18,996 unique mutations, collected from 587,357 human superficial melanoma’s gene samples.

Figure 2A shows that the EFD is an unimodal, skewed and has a heavy tail on the right that is in the log-log plot. The mean value of the number of unique mutated gene samples consists of 30.9, median 20, mode 5. One mutation per gene was observed in 320 samples, a maximum number of unique mutated samples per gene equals 626.

**Figure 2:**
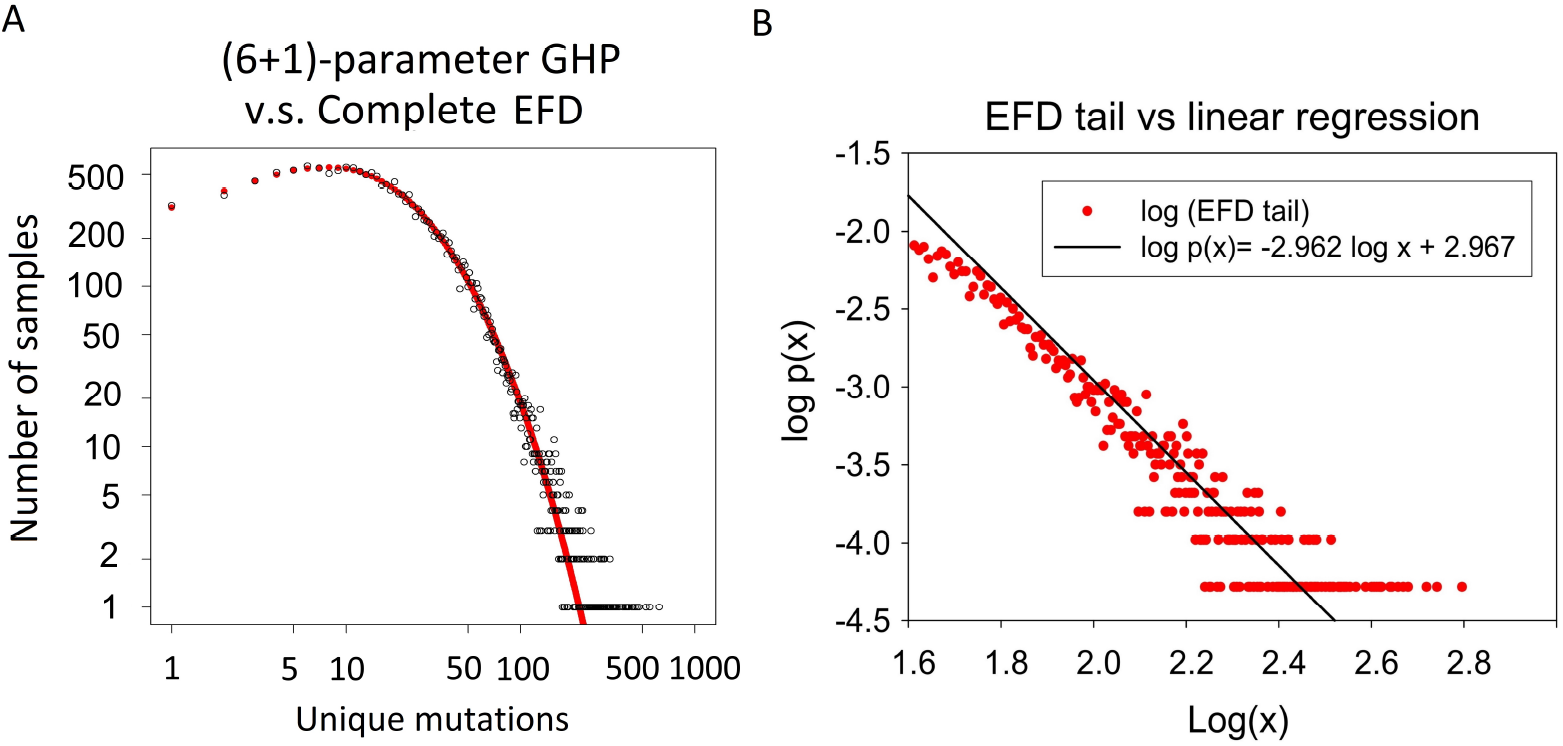
Frequency distribution of superficial m elanoma mutations. 7-parameter Generalized PDF. A: Example of stationary BDP-derived PDFs model’s parameterization using complex form of the EFD. The frequency distribution of the number of unique mutated samples of gene loci detected in the large collection of human melanomas have modeled by (6+1)-parameter GHP distribution. Estimated parameters and goodness of fit criteria values are 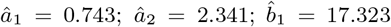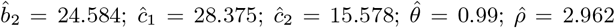 Ψ = 5.641. B: Regression model with negative slope 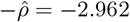 approximates asymptotic decreasing the EFD tail. Correlation coefficient between experimental and predicted data is 0.874; *p* – *value* < 0.0001 (t-test, F-test). The EFD r.v. *x* interval [65-625] is shown on axis *X* on log_10_ scale.

We parameterized the (3d+1)-GHP (27) with 7 parameters. We utilized the bounded quasi-Newton limited-memory BFGS (L-BFGS-B) algorithm to estimate the parameters, using our goodness of fit Ψ criteria [26, 28].

We were able to parameterize our (6+1)-parameter PDF using (27)at a high level of the goodness of fit criteria value (Figure 2 A). Furthermore, our results show that the parameterized PDF fits well not only to the long tail of the EFD but also to the entire random variable domain of the EFD (Figure 2A). Seven estimated parameter values, calculated 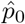 and the Ψ criteria value are showed on the legend of Figure 2.

We observed that the estimated parameter 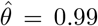 and 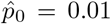. According to these estimates, we suggest (see Discussion) that we could use the parameters to analyze the Theorem 1 and Remark 1 predictions referring to parameter *ρ* - the power law exponent at asymptotic values of *x* (observed number of unique melanoma somatic mutations in gene samples). According to the estimated values of our model parameters and Theorem 1 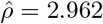. Figure 2B shows that the linear regression function with a negative slope equals −2.962 fits to the experimentally observed tail end of the EFD for the r.v. values larger then 115 unique gene loci mutated samples.

## 6 Discussion and Conclusion

In this study, we introduced and characterized a multi-parameter family of generalized GHP models, generated by the stationary g-BDP and defined by PMF (27) and associated probability measure called (3d+1)-GHP (28). We specified a generalized probability subfamily called the 3d-RGHP as a regularly varying model associated with PMF (38).

Our 3d-RGHP model represents a broad family of regularly varying PDFs with long right tails. To offer 3 d-RGHP as a *new* EFD in bioinformatics, the *statistical facts*, such as regular variation at infinity, the exhibition of asymptotically constant slowly varying component, upward/downward convexity, unimodality, and convexity, were established. The non-linear dependence of the transition rates from a state of the stationary BDP allows developing many applications. In particular, the model presents a rigorous and flexible analytical framework for analysis of the skewed distribution functions generated by BDP observed in big data bioinformatics.

The number of unique mutated samples in a gene of melanoma genome shows a skewed EFD distribution with a long right tail. Skewness is strongly characterized by values and order of mode=5, median=20 and mean=31. This proposes that the mutagenesis in melanoma cells is a highly-intensive and driven multiple probabilistic mechanisms. Our model successfully describes the complexity of the EFD and the results of our estimates (Figure 2A) that demonstrated the ability of model (27) to quantify statistical characteristics of the EFD in the contexts of the skewed long tail and its potential power law properties.

Meanwhile, according to our results, a probability of non-observed somatic mutation consists of only 1.1% suggesting that the experimental outcome space is almost fully detected. Prediction of the *p*_0_ estimation provides a statistical basis for unbiased PDF identification.

Importantly, Figure 2A showed that our (6+1)-GHP PDF accurately fits the EFD on the entire random variable domain *X* = *x*; *x* ∈ [1, 625]. Assuming that birth and death distributions are stationary processes, we noted that mutagenesis in the melanoma genome has an extreme (near ‘critical’) regime defined by the value of 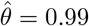, that according to our theory is corresponding to regularly varying transition rates *θ* = 1. Furthermore, by our estimation, the experimentally observed power law tail end exponent equals 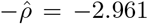. Figure 2B shows that the linear regression with negative slope equals −2.962 fits well a segment of r.v. *x* at the tail end of our EFD supporting our Theorem 1 and Remark 1. These results suggest that our multi-parameter probabilistic models may be practically used for real bioinformatics and other big data analysis.

Interestingly, these results suggest that the probability values of the power law exponent asymptotic are reduced significantly faster at *x* → ∞ in comparison to the standard Zipf law, having a slower decay exponent parameter (−*ρ* ≈ − 2). This value is commonly observed in many EFDs with a power law-like tail. Thus, even the EFD has a long skewed tail region on the right, the end of the tail of the region with asymptotic power law property could be relatively short. Figures 2(A-B) showed that the right side tail of our parameterized PDF monotonically deviates from the direct line when *x*-value increases. The EFD shows the region with power-law properly is overdispersed. A possible explanation of the reduction of a power-law property region is the non-linear dependence of the ratio of birth and death rate functions from state value.

Figures 1(A-D) showed the PMFs at used parameter sets are unimodal with mode value 1. Numerical analysis showed that the length and shape of the right-side tails are significantly varied with parameter value changes. However, the PMF tail’s length and shape are sensitive to the parameter values. Figures 1(E-H) demonstrated the log-log plot of the double-truncated distribution representing the power law tails of the PDF presented in Figures 1(A-D) respectively. For each set of the parameters, the cut-off value interval was defined according to the linearity of the power law exponents of the tails in the log-log plot. The log-log plots of Figures 1(E-H) showed that on relatively long intervals of random variable on the right-side of the 3d-RGHP, the power law-like tail can be significantly deviated from the straight line.

Thus, our numerical analysis of the 3d-RGHP and an example of the real data parameterization of the model suggest that in dependent of the parameter value sets, the right-side tail of the PMF could be increased or decreased however, power law-like exponent asymptotic may be approached at very large r.v. values at the tail end; such interval could be relatively short, unstable and sensitive to sample size.

In these contexts, we point out that the concept of scale-free distribution [8] model and its multiple variants, that focus on the power law tail properties of the distribution, ignore the experiment outcome space completeness and the non-linear and multiple probabilistic mechanisms, that makes such theories quite restricted, non-rigorous and practically non-applicable [10, 21, 25, 26, 27, 28, 29, 30, 31, 32, 38, 39].

Our approach to modeling of the stochastic process considering complexity, non-linear interactions/connectivity, integrity, completeness, birth and death mechanisms, stationary and transitions allows deep and flexible theory development and provides a natural bridge between modeling, algorithms and statistical methods development and the real-life applications needs. Importantly, our theory could be used for statistical hypotheses testing in big data analysis, and our model estimates and predictions are testable.

Big data sets exist and are rapidly collected not only in bioinformatics and biomedical sciences but also in all kinds of data-intensive areas corresponding to modern physics, astronomy, material sciences, geophysics, internet, technology, epidemiology, economics, finance, social sciences, community networks, visitors monitoring, etc. Skewed distributions of complex form and extreme characteristics regularly occur in data sets from these disciplines. We suggest that our g-BDP models may be useful in many such cases.

In summary:

We suggest that the many observed skewed discrete distributions are generated by non-linear and multiple independent stochastic process and their combinations;

Our 3d-RGHP model presents a flexible analytical framework for analysis of the skewed distribution functions generated by BDP observed in bioinformatics and big data science;

Our results suggest that the 3d-RGHP model can be used for statistical analysis and identification of probabilistic mechanisms and significant characteristics of evolving systems whose EFD exhibits complex form and a heavy tail at critical regime;

Our model could help better understand stochastic processes driving evolution of biological complexity and the pathological role of extreme values.

